# Cell fitness is an omniphenotype

**DOI:** 10.1101/487157

**Authors:** Nicholas C. Jacobs, Jiwoong Park, Nancy L. Saccone, Timothy R. Peterson

**Affiliations:** Department of Medicine, Division of Basic and Biomedical Sciences, Washington University School of Medicine, 660 S. Euclid Ave., St. Louis, MO 63110, USA; Computational and Systems Biology, Washington University School of Medicine, 660 S. Euclid Ave., St. Louis, MO 63110, USA; Molecular and Cellular Biology programs, Washington University School of Medicine, 660 S. Euclid Ave., St. Louis, MO 63110, USA; Department of Genetics, Washington University School of Medicine, 660 S. Euclid Ave., St. Louis, MO 63110, USA; Division of Biostatistics, Washington University School of Medicine, 660 S. Euclid Ave., St. Louis, MO 63110, USA; Institute for Public Health, Washington University School of Medicine, 660 S. Euclid Ave., St. Louis, MO 63110, USA; BIOIO, 4340 Duncan Ave., Suite 236, St. Louis, MO 63110, USA; Healthspan Technologies, Inc., 4340 Duncan Ave. Suite 265, St. Louis, MO 63110, USA

## Abstract

Moore’s law states that computers get faster and less expensive over time. In contrast in biopharma, there is the reverse spelling, Eroom’s law, which states that drug discovery is getting slower and costing more money every year. Herein, we propose a solution to this problem. We put forth a consensus algorithm for inexpensively and rapidly prioritizing new factors of interest (e.g., a gene or drug) in human disease research. Specifically, we argue for synthetic interaction testing in mammalian cells using cell fitness – which reflect changes in cell number that could be due many effects – as a readout to judge the potential of the new factor. That is, if we combine perturbing a known factor with perturbing an unknown factor and they produce a synergistic, i.e., multiplicative rather than additive cell fitness phenotype, this justifies proceeding with the unknown gene/drug in more complex models where the known perturbation is already validated. This recommendation is backed by the following evidence we demonstrate herein: 1) human genes currently known to be important to cell fitness involve nearly all classifications of cellular and molecular processes; 2) Nearly all human genes important in cancer – a disease defined by altered cell number – are also important in other common diseases; 3) Many drugs affect a patient’s condition and the fitness of their cells comparably. We provide proof of concept of the Omniphenotype model using the widely used osteoporosis drug, bisphosphonates, implicating its mechanism of action (MoA) genes, ATRAID, SLC37A3, and FDPS, as potential gerotargets for neurodegenerative conditions. Taken together, these findings suggest cell fitness could be a broadly applicable phenotype for understanding gene, disease, and drug function. Measuring cell fitness is robust and requires little time and money. These are features that have long been capitalized on by pioneers using model organisms that we hope more mammalian biologists will recognize.

**Short summary:** Cell fitness is a biological hash function that enables interoperability of biomedical data.

## INTRODUCTION

Despite or maybe because of technology improvements there is currently no solution for Eroom’s law. Eroom’s law is the observation made by Scannell et al. that drug development gets slower and more expensive over time (1, 2). It might be said that this problem is one of lack of consensus: There are so many approaches to solve biomedical problems, and a growing number of approaches, that the lack of coordination around the most high-yield approaches increases the time and money spent. To address this, perhaps we need to look to lessons on how other fields think about problems of coordination. Consider how the longstanding computer science problem known as the Byzantine Generals Problem was solved. The Byzantine Generals Problem was articulated by computer scientists working for NASA back in the 1970’s to think through air traffic control risks considering the realization that computers were soon going to be flying planes (3, 4). The general form of the Byzantine Generals Problem describes a leaderless situation where involved parties must agree on a single strategy in order to avoid failure, but where some of the involved parties are unreliable or are disseminating unreliable information (5). Interestingly, like with important problems in biology that were solved using knowledge from other disciplines, such as with physics in helping solve the structure of DNA, key insights to the Byzantine Generals Problem came from outside of computer science. Namely, in economics it was solved by a consensus algorithm based on “proof-of-work” that is now used to run the Bitcoin peer-to-peer monetary system (6). Developing such a consensus experimental approach to optimize biomedical research does not exist currently. Needless to say, such an approach would be extremely valuable if one could be identified.

Consensus in biomedical research has historically taken the form of researchers agreeing on what biological model systems they will focus their labs around. Interestingly, relatively simple model systems such as yeast and worms have made a disproportionate contribution to human health (7–9). For example, the broadly medically important mechanistic target of rapamycin (mTOR) pathway was first discovered in yeast using a cell fitness-based, i.e., cell counting-based, screen (10, 11). Moreover, evidence shows that even more complex models such as mice aren’t representative models of human diseases (12, 13). Some researchers are nevertheless pushing forward and focusing on creating new ways to accelerate human disease research using model organisms (14–16). Yet, many important human genes aren’t conserved in model organisms. Moreover, even if one uses a mammalian system, it is assumed that the context being studied, e.g., tissue type, must be taken into an account. There is therefore a need for models that strike a balance between the ease and robustness of model organisms while preserving the signaling connectivity important to human disease.

Modeling disease has gained new relevance as the number of human population genetic studies has exploded (17). Thousands of genes are being implicated in human diseases across categories from cancer to autism, Alzheimer’s, and diabetes. Among the surprises from these data are findings such as “cancer” genes – genes understood to be important in cancer – being linked to diseases seemingly unrelated to cancer. For example, one of the earliest genes found to be mutated in the behavioral condition, autism (18), is the well-known tumor suppressor gene, PTEN. On first pass, it might be difficult to glean what autism might have to do with the uncontrolled cell proliferation that PTEN inactivation causes. However, it is clearer when one considers that patients with autism often have larger brains and more cortical cells (19). Nevertheless, the extent to which the various functions of genes like PTEN are relevant to disparate diseases begs the questions of how we molecularly classify and model disease and whether we truly understand the function of these genes beyond cancer.

As with disease genes, new treatments have often suffered from biases that prevented us from seeing their value outside of the context in which they are originally characterized. For example, cholesterol-lowering statins were initially held back from the clinic because they were shown to be toxic in animal models. Those early findings scared off pharmaceutical companies because they suggested statins might not be safe for humans (20, 21). Merck eventually realized this toxicity was on-target related to the drug mechanism of action and not generic off-target activity, and the statins went on to be one of the biggest success stories in drug development history (22). The successes of drug “repositioning” also reveal to us that drugs can function in multiple “unrelated” contexts. For example, the malarial drug, hydroxychloroquine (Plaquenil), is now routinely used to treat rheumatoid arthritis and autoimmune disorders (23). There is also chemotherapeutics being considered for depression (24). Drug repositioning makes sense from a molecular perspective because many drug side effects could actually be on-target effects (25). To take it further, now with so much knowledge in the “-omic” era, it might be the rule rather than the exception that a mechanistic target for a drug has many functions in the body. For example, the serotonin transporter, SLC6A4 (a.k.a. SERT), is the target of the widely prescribed antidepressants, Prozac and Zoloft. SLC6A4 is expressed at high levels in the intestines and lungs (26), and regulates gut motility and pulmonary blood flow (27, 28) in addition to mood. This demonstrates the need to find a better way to understand the diverse roles of drugs and their targets.

Herein, we address these aforementioned issues. First, we quantify costs incurred for studying proteins for eventual drug development and demonstrate that continuing with the status quo will require prohibitive amounts of time and money. Then, we present evidence that synthetic interaction testing using cell fitness provides a novel solution to this challenge. Specifically, by leveraging diverse data types: gene-inactivation screens in cells, PubMed, and human genome-wide association studies (GWAS), we demonstrate that cell fitness-based synthetic interaction testing would be predicted to yield important insight on a new factor of interest (e.g., gene or drug) irrespective of the ultimate phenotype of interest for that factor. This predictive power comes from two related findings. One, that cell fitness is altered upon perturbation of most, if not all, biological pathways. Two, that a gene’s or drug’s relevance to a disease is correlated with its relevance to an in vivo proxy of cell fitness, cancer.

Based on these findings, we propose that synthetic interaction testing in mammalian cells using cell number/fitness as a readout is a relevant and scalable approach to understand surprisingly diverse types of molecular functions, diseases, and treatments. This proposal requires a change in thinking because synthetic interaction testing using human cells has historically almost always been restricted to the context of cancer (29, 30). This makes sense considering cancer has become defined by alterations in individual phenotypes that contribute to cell number/fitness, e.g., proliferation and apoptosis (31, 32). However, we argue fitness-based synthetic interaction testing has a much more general purpose. Our evidence indicates that cell fitness is a generalizable phenotype because it is an aggregation of individual phenotypes. To the extent that it might be an aggregation of all possible phenotypes – an omniphenotype – suggests its potential as a pan-disease model for biological discovery and drug development.

## RESULTS

### Problem: The current approach to studying human disease biology is time-consuming and costly

Proteins are key building blocks of life. They are also what most drugs are designed to target. Proteins interact with each other to perform complex functions, such as DNA replication, ATP production, and the trafficking and degradation of various molecules. The human protein interactome is vast, and by one estimate 8.8M of them could be essential for viability (33). We asked how much time and money has been spent on studying these interactions to date.

When we look at a high-confidence list of 21,729 protein-protein interactions (PPIs) (34), NIH funded grants have studied 3.3% of these pairs (717 pairs) over the past 20 years and $3.5B has been spent on them. At the current rate, it can be estimated to take many more decades even centuries and costing hundreds of billions or even trillions to study all 21,729 interactions. Notably, this does not include investment to develop drugs based on this information, which is estimated to push the numbers up potentially up to 66X more (35) (Fig. 1A). Also, these numbers are for the 21,729 pairs, which is less than 1% of the total 8.8M potentially important ones. While these estimates do not factor in improvements in data analysis, e.g., artificial intelligence (AI), and PPIs are only a subset of the problem space, clearly, we could make improvements to understand human genes in a more time- and cost-effective manner.

**Figure 1.**
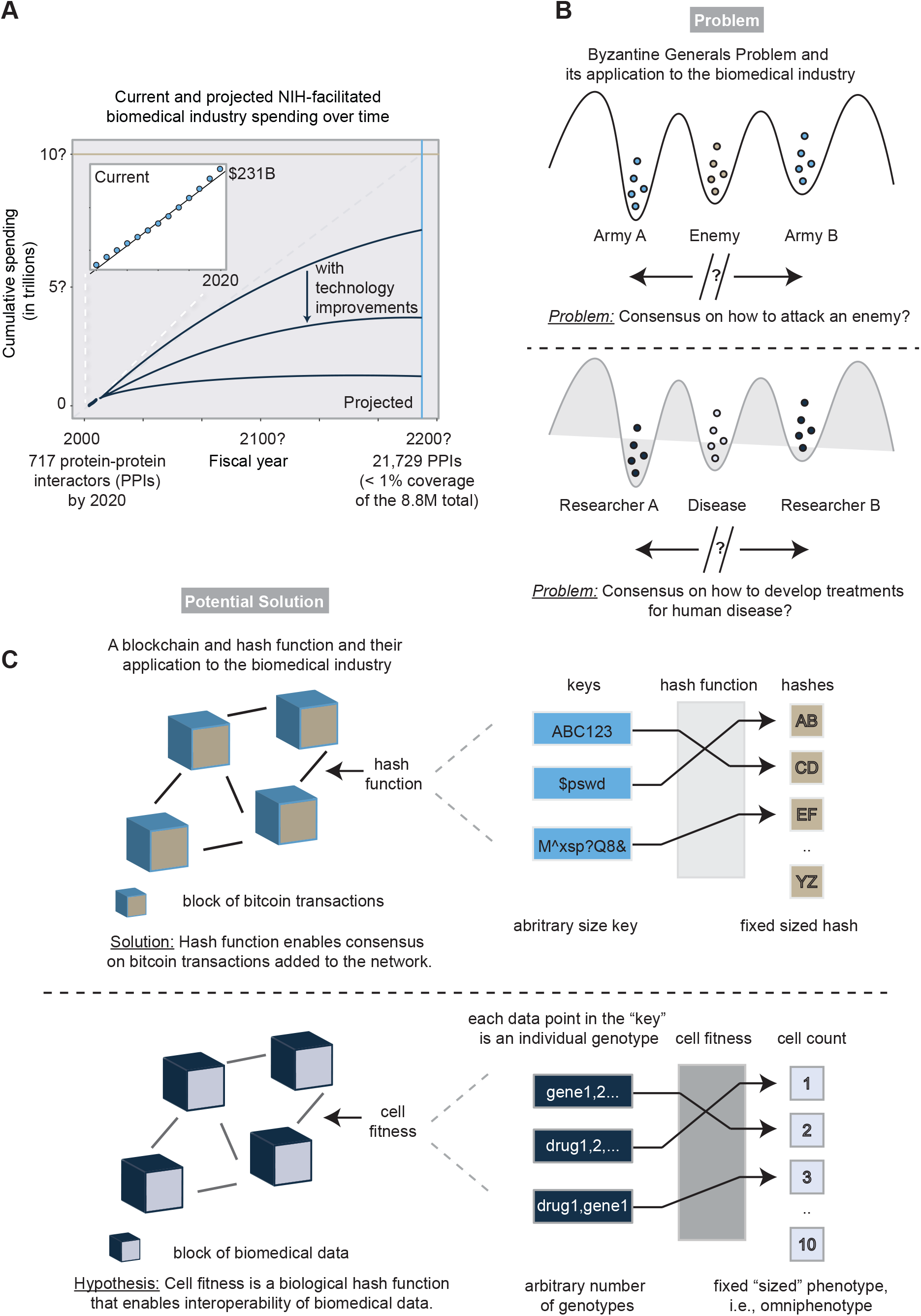
The problems of and a potential solution for the lack of consensus-building mechanisms in biomedical research. **(A)** Current and projected time and money spending estimates to develop drugs for high-confidence protein-protein interaction (PPI) pairs obtained from Huttlin et al. (34, 71). The projected budget is a line of best fit based on the cumulative NIH grant money spent on studying gene pairs multiplied by how much is needed to convert this funding to FDA approved drugs, i.e., 44,200/670 = 66X (35). The inset graph represents the current spending estimate that is extrapolated to make the projected spending and time estimates. The intersection of the light blue and tan lines represent the projected date and costs, respectively, when all drugs will be identified for each PPI. **(B)** A schematic of the Byzantine Generals Problem and its application to the biomedical industry. In a real-world scenario, achieving consensus becomes much more difficult when there are many parties involved (greater than the two depicted) that are not working together. **(C)** Data blocks are chained together using a robust algorithm called a hash function to create consensus understanding. In the case of bitcoin, the consensus determines who owns what bitcoin. To make an analogy on how the concept of a blockchain can address Eroom’s law, we hypothesize that cell fitness is a biological hash function that enables interoperability of biomedical data from different research efforts.

We considered the biomedical industry as a poorly coordinating network. To make the network more efficient, we considered lessons from computer science and the study of distributed systems. A famous problem in distributed systems that seemed relevant to Eroom’s law is the Byzantine Generals Problem. The Byzantine Generals Problem is a hypothetical situation in which a group of generals lead armies that are geographically separated from each other (such as on other sides of two hills), which makes their communication difficult (Fig. 1B). They must choose either to attack the enemy (which is in the valley between them) or retreat. A half-hearted attempt at either will result in a rout, and so the generals must come to a consensus decision on which approach to take. To extend the analogy to the biomedical industry, we considered how the lack of consensus by researchers on what experiments should be prioritized has led to an inability to win the “war on cancer” as well as other ‘battles’ against other diseases (Fig. 1B) (36).

### A Potential Solution: Synthetic interaction testing using cell fitness

At the heart of the Bitcoin’s solution to the Byzantine General’s Problem is its blockchain. A blockchain is a publicly viewable and editable database (Fig. 1C). It has comparisons to a collection of Google or Excel spreadsheets in that everyone anywhere can read and write to them. However, a key distinction is that on a blockchain one can’t overwrite other people’s additions. This cryptographically secured step maintains the fidelity of all the data, which thus leads to a universally shared understanding of it. Consensus in a blockchain is achieved using hash functions, which transform arbitrary sized data into fixed sized hashes (Fig. 1C). The point of hashes is that they are much easier to analyze relative to the data that comprises them. This allows people to relatively easily trust that the data as a whole is accurate without having to verify each individual piece of it themselves. Researchers are starting to articulate how biomedical research could be aided by the concept of blockchains (37–39). The volume of data in biomedical research is exploding, yet it is growing rapidly in different silos. We wondered what kind of biological ‘hash’ function might be well suited to faithfully ‘chain’ together disparate ‘blocks’ of biomedical data. Recently, the Cancer Dependency Map, a.k.a., DepMap, shed important light on PPIs across the human proteome (40, 41). This is interesting because the DepMap only required cell fitness as a phenotype to identify the PPIs. Another study, PRISM, used a similar approach as DepMap, but instead of mutating genes to perturb cell fitness, it used drugs (42). Both DepMap and PRISM relied on many of the same cell lines to make their predictions. Thus, we asked if one were to ‘chain’ together the DepMap and PRISM ‘blocks’ of cell fitness data, how many drugs targeting the aforementioned 21,729 PPIs might be identified. We estimate that these two datasets alone could provide a first pass at drugging 27% of these PPIs, representing $2.67T and 149 years of future money and time spent if they weren’t used (Supp. Table 1). The underlying hash function-like, consensus algorithm we hypothesize one would use to ‘chain’ together the DepMap and PRISM involves imputing synthetic interactions between genes and drugs from their cell fitness data (Fig. 1C). Therefore, to explain how we arrived at this 27% number and how we might eventually get to 100% coverage using cell fitness data, it is first important to explain synthetic interaction testing.

A simple way to understand synthetic interactions is to consider two genes. It is said there is a synthetic interaction between two genes if the combined effect of mutating both on cell fitness, hereafter denoted by ”*ρ*” (as in relative cell fitness or phenotype), is much greater than the effects of the individual mutations (Fig. 2). If there is an interaction, it can be either be a negative synergy (Fig. 2, 5^th^ from left panel), where the double mutant cells grow much worse (a.k.a., synthetic lethality), or a positive synergy, where the cells grow much better (a.k.a., synthetic buffering), respectively, than the individual mutant cells (Fig. 2, 6^th^ from left panel). Both phenotypes demonstrate a synergy on a genetic level between these two genes).

**Figure 2.**
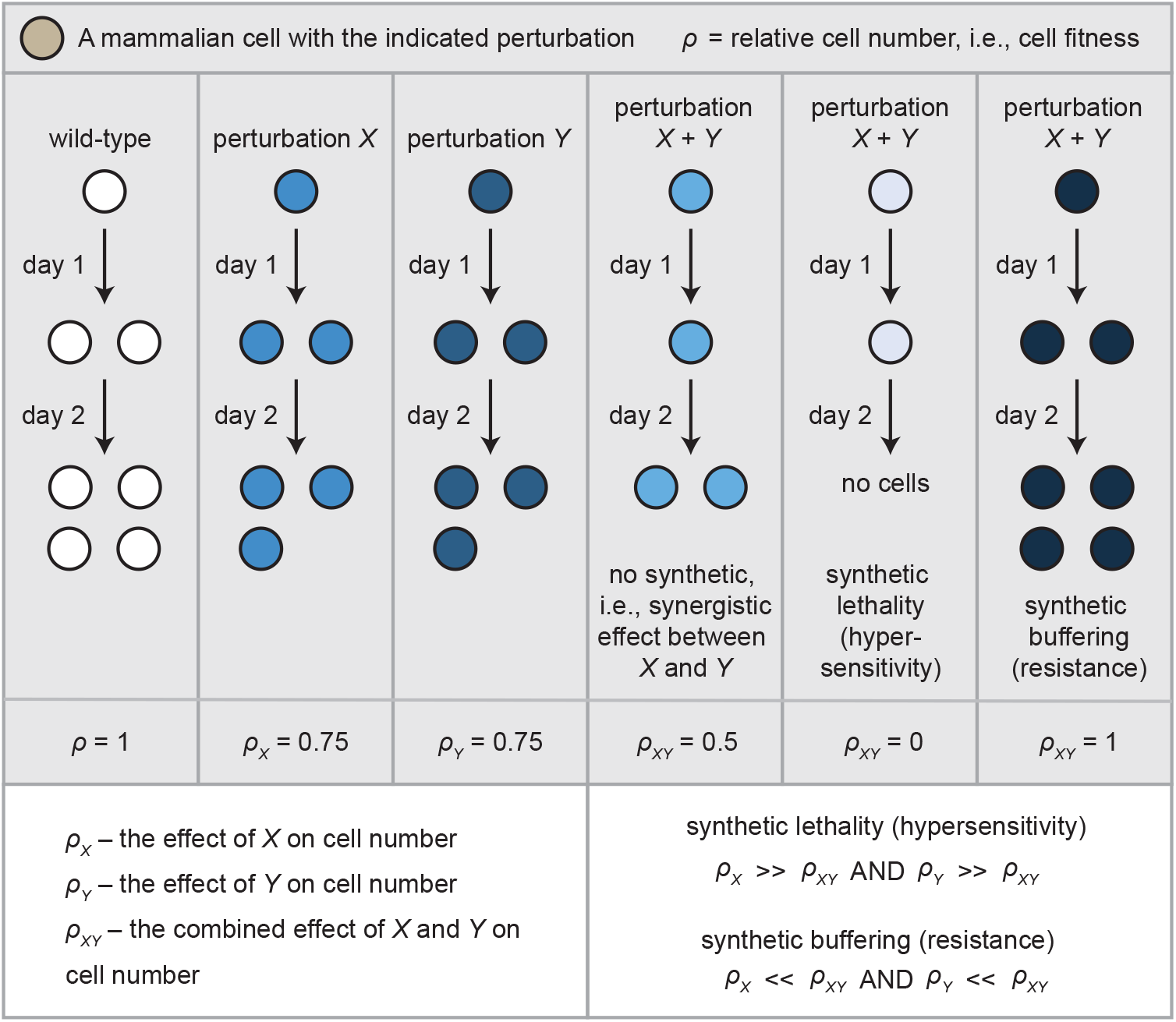
A current, commonly used framework for synthetic interaction testing. A synthetic interaction between two perturbations, *X* and *Y*, means the effect of *X* combined with *Y* on cell fitness significantly differs from the expected effect based on each effect alone; typically, an additive effect is expected. Synthetic interactions can cause lethality or hypersensitivity, or can be buffering or confer resistance. Both are depicted in the figure where *ρ* – the phenotype – is relative cell number. Synthetic effects can be thought of as multiplicative or exponential rather than linear or additive, as is the case in the condition (4^th^ column from left) where 1 - *ρ*_x_ (0.75) + 1 - *ρ*_y_ (0.75) = *ρ*_xy_ = 0.5. Note: In some scenarios, a perturbation is lethal on its own, which precludes an investigation of its synthetic lethal but not its buffering interactions. On the other hand, the lack of a perturbation’s effect on its own on cell fitness does not preclude one from determining if it has a synthetic interaction with another perturbation.

Amongst the ~400 million possible synthetic interactions between protein coding human genes, less than 0.1% have been mapped to date (43). It is challenging to scale synthetic interaction testing in human cells both because of the size of the experiments needed and because sometimes neither interactor has a known function in the ultimate biological context of interest (44). Therefore, we sought to develop a framework that would allow us to infer synthetic interactions at scale as well as leverage interactors of known significance to explain interactors of unknown significance. The convention of characterizing an unknown in the context of a known is a typical approach taken by biologists for making discoveries. To formalize our approach, we hereafter refer to the known interactor in a synthetic interaction pair as *K,* and the unknown as *U.* With this convention, *U* is what’s new and interests the researcher, whereas *K* is a factor that’s known in the field. *U* and *K* could each represent a gene, disease, or drug. Throughout this work, we provide three pieces of evidence that taken together provide a generalizable framework for using cell fitness-based synthetic interaction testing to solve for *U* using *K*.

### Evidence #1: Nearly all cell processes affect cell fitness

Recent works by Hart et al., Blomen et al., and Wang et al. identified “cell essential” genes – genes that affect cell fitness to the degree that they are required for cell viability (45–47) – and we wondered what GO pathways they participate in. In this case, we took the above published lists of cell essential genes and considered the synthetic interaction testing as follows: the known, *K,* is the gene that is essential in at least one cell type and the unknown, *U,* is the synthetically-interacting GO pathway of interest. GO pathways are collections of genes. As precedence for this approach of considering *U* as a collection of genes rather than an individual gene we applied the principal of the minimal cut set (MCS). MCS is an engineering term used by biologists to mean the minimal number of deficiencies that would prevent a signaling network from functioning (48, 49). MCS fits well with the concept of synthetic lethality (29), and this was recognized by Francisco Planes and colleagues to identify RRM1 as an essential gene in certain multiple myeloma cell lines (48). MCS also relates to the concept of co-dependency where two genes are said to be co-dependent if their gene deficiency cell fitness profiles correlate with each other across cell lines (50).

We generalized the idea of MCS to hypothesize that different cell types might be relatively defective in components of specific GO pathways such that if gene(s) in one of these pathways were knocked out, they would produce a synthetic lethal interaction with those components (For clarity throughout this study, by “genes”, we refer to human protein coding genes). Applying this MCS approach, we found the essential genes from the Hart, Blomen, and Wang studies are involved in diverse GO processes and the number of GO processes they participate in scales with the number of cell lines examined (Fig. 3A, Supp. Table 1). For example, if one cell line is assessed, at most 2054 out of 21246 total genes assessed were essential and these essential genes involve at most 36% of all possible GO categories (6444 out of 17793. 17793 represents the number of unique – i.e., irrespective of the GO hierarchy - human gene-associated GO categories). In examining five cell lines, 3916 essential genes were identified that involve 54% of all GO categories (9577/17793). With ten cell lines, 6332 essential genes were identified that involve 68% of all GO categories (12067/17793). Therefore, essential genes involve the majority of GO processes. Because the Fig. 3A graph appears to approach 1, this suggests if enough cell types are tested, potentially all genes and GO processes would be essential in at least some cell context. Consistent with this, when we relaxed the significance on the cell fitness scores from p < 0.05 to p < 0.10, this relaxed threshold encompasses nearly all GO processes and genes (Fig. 3A, 97.2%, 17291/17793 GO terms; 17473/21246 genes). Taken together, this suggests the majority if not all GO processes and their associated genes affect cell fitness.

**Figure 3.**
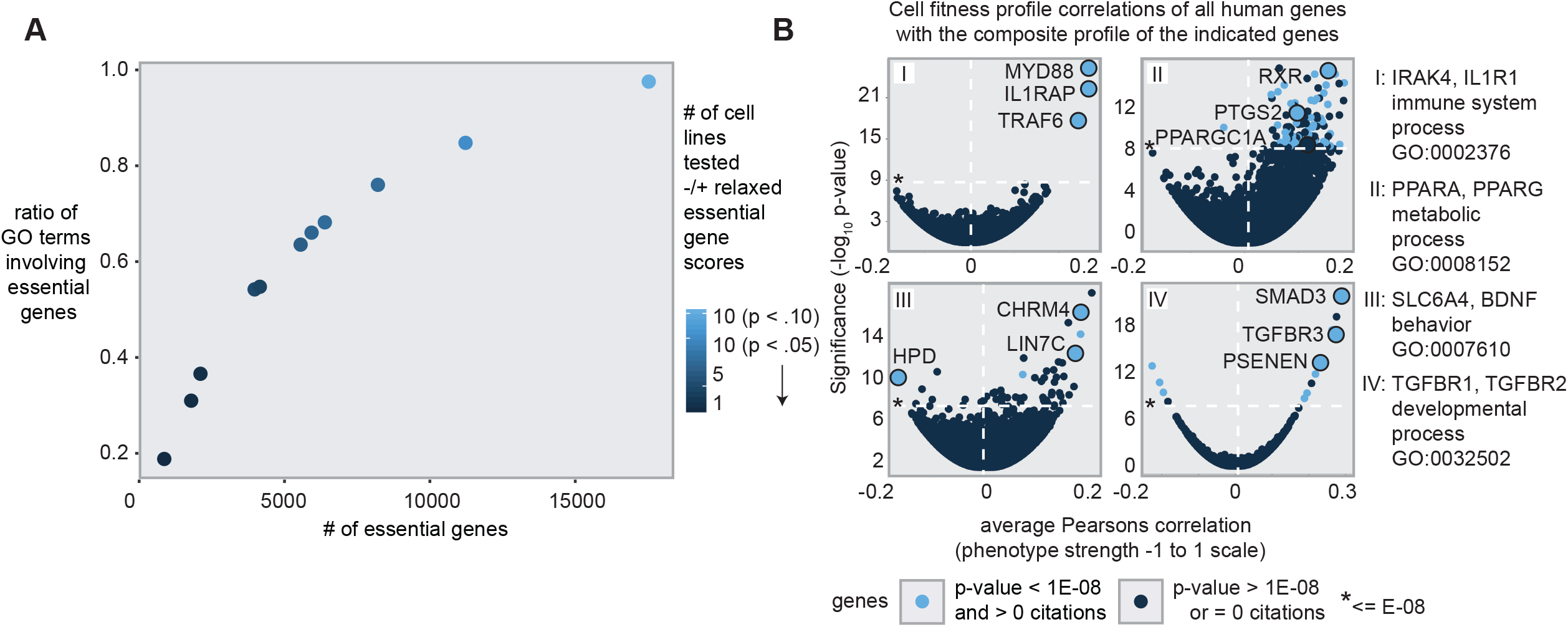
Nearly all GO processes are essential for cell fitness. **(A)** The y-axis is the ratio of the number of distinct GO terms linked to the essential genes in the tested cell lines divided by the total number of distinct GO terms (17,793) for all tested human genes (21,246). The x-axis is the number of essential genes determined for the indicated number of cell lines. “Relaxed” means p < 0.10 (as opposed to p < 0.05) cell fitness scoring. **(B)** Fitness profile correlations across 563 human cell lines of mutant cells of genes from the same GO categories. The statistically significant genes that have been most highly co-cited with the indicated genes are color-coded light blue. P-values were false discovery rate (FDR) corrected using the Benjamini and Hochberg method (72).

Recently, genome-scale gene inactivation screening has been performed on hundreds of human cell lines (40). Using this data we reasoned that the single gene mutant cell fitness profiles for genes in the same GO processes would strongly correlate across cell lines. We selected gene pairs from diverse GO categories (immune system process (GO:0002376), behavior (GO:0007610), metabolic process (GO:0008152), and developmental process (GO:0032502) and asked how the fitness profiles of other members of the same GO category would rank compared with all 17,634 fitness profiles that were assessed. These categories were chosen because they are direct children of the top-level “Biological Process” GO category (GO:0008150, with a “is_a” relationship) that cover broad areas of human health and have similar numbers of genes associated with them. Impressively, highly co-cited genes from each top-level GO category were highly correlated in their fitness profiles (p < 1e-8, Fig. 3B, Supp. Table 1). Gene co-citation is a human determined connection between two genes whereas cell fitness correlations are machine determined. Therefore, that all significantly correlating cell fitness profiles were also enriched as being co-cited (compare light and dark blue in Fig. 3B, hypergeometric distribution model enrichment p-values: immune system process p = 1.4E-4; metabolic process p = 6.9E-3; development p = 2.9E-3; behavior p = 2.8E-2) suggests purely computational approaches using cell fitness can accurately predict gene function for disparate biological processes.

### Evidence #2: Nearly all common disease genes are cancer genes

The omnigenic model states that all genes expressed in disease relevant tissues are important to the disease (51). Yet, there is a scale and some genes are more important than others. These relatively more important genes are referred to as “core” (a.k.a. “hub”) genes. We reasoned that the published literature with its millions of peer reviewed experiments would be a comprehensive way to inform us on which genes are core genes for a given common disease. At the same time, we were interested in leveraging the literature to identify genes that would most strongly synthetically interact with these core genes.

Here we defined a synthetic interaction as follows: the gene(s) mentioned in a citation is the unknown, *U*. In this convention we make the simplifying assumption that this gene(s) is not known in whatever context was studied, e.g., diabetes, which is why it warrants a publication. Whereas all other already published genes in that context at the time of publication we consider as the known, *K*. To measure synthetic interaction strength between *U* and *K*, we assessed their co-occurrence with the term, “cancer” as a proxy for cell fitness because cancer is a common disease defined by altered cell number. Specifically, we assessed all citations linked to the MeSH term “neoplasm” vs. other MeSH terms in the Disease category (we hereafter refer to these as “cancer” vs. “non-cancer” citations, see Methods for more details). For context, the non-cancer literature includes conditions like the top ten most common causes of death according to the World Health Organization (52) and other sources: Alzheimer’s, cardiovascular disease, diabetes, obesity, depression, infection, osteoporosis, hypertension, stroke, and inflammation. In analyzing all currently available PubMed citations (~30M), we identified a compelling relationship (Correlation coefficient, r = 0.624): most genes are cited in the cancer literature to a similar extent as with the non-cancer literature (Fig. 4A, Supp. Table 1).

**Figure 4.**
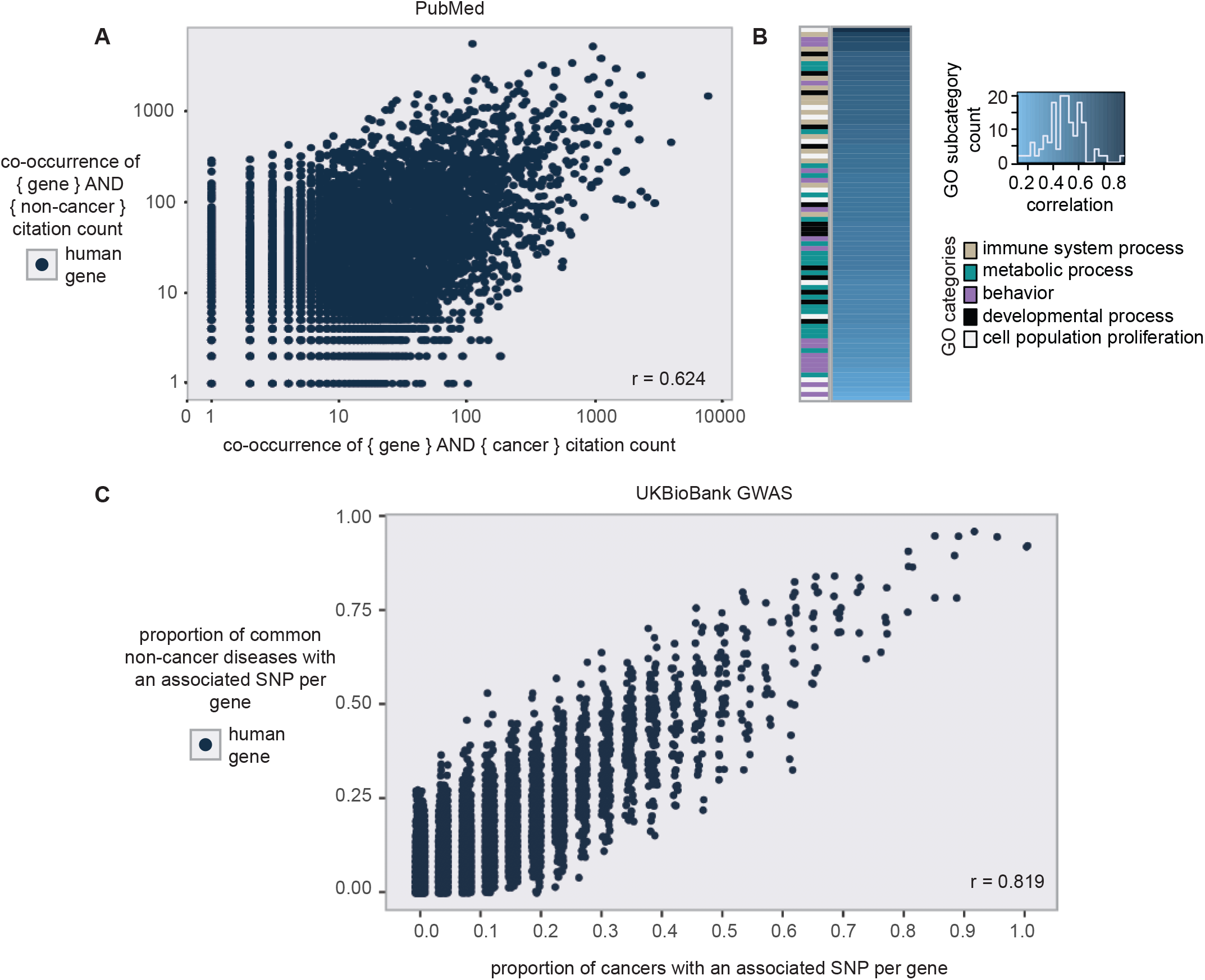
Nearly all common disease genes are cancer genes. **(A)** The co-occurrences of each human gene with cancer (x-axis) and non-cancer (y-axis) conditions in all currently existing ~30 million PubMed citations were counted. The non-cancer conditions include all MeSH terms in the Disease category not classified with the “neoplasm” MeSH classifier. It includes common conditions like infection, Alzheimer’s, cardiovascular disease, diabetes, obesity, depression, inflammation, osteoporosis, hypertension, and stroke amongst many other MeSH classifiers. Line of best fit - Slope, 0.67; Correlation coefficient, r = 0.624. **(B)** Gene ontology (GO) classifications that comprise the non-cancer/cancer correlation in (A). Categories are sorted from dark blue representing a stronger correlating GO category to lighter blue representing a weaker correlating category. **(C)** The proportion of non-cancer diseases vs. cancer diseases that have an associated SNP (p-value < .001) in UKBioBank for each protein-coding human gene. These proportions are out of the top 100 conditions, 26 cancer diseases and 74 non-cancer diseases, based on co-citation with genes. Correlation coefficient, r = .819.

Some non-cancer citations such as those that focus on a gene’s physiological function or those that focus on rare diseases could undermine our goal of making an apples-to-apples comparison of non-cancer diseases with cancer. To more strictly determine whether the cancer vs. non-cancer correlation pertains to common diseases, we restricted the non-cancer set to common conditions that include the top causes of death according to the World Health Organization (52) and other sources: Alzheimer’s, cardiovascular disease, diabetes, obesity, depression, infection, osteoporosis, hypertension, stroke, and inflammation (~10M citations). This more stringent analysis surprisingly produced a stronger correlation (r = 0.69) than the more all-encompassing non-cancer citations list (Fig. S1, Supp. Table 1). These strong correlations suggest for many non-cancer diseases, genes important to them that are not yet known can be identified using synthetic interaction testing. Also, that some genes are cited much more than others across disease categories is consistent with the multiple functions of core/hub genes.

We analyzed the pattern we detected in Fig. 4A in more detail. To understand what GO categories led to better cancer/non-cancer citation correlations, we binned the cancer and non-cancer citations for all genes by their top-level GO categories we used in Fig. 3: immune system process, behavior, metabolic process, and developmental process. We also included cell population proliferation as that is directly related to cancer. We found no clear pattern of overtly cancer-related GO sub-categories that contribute most vs. least to the high cancer/non-cancer citation correlation (Fig. 4B, Supp. Table 1). For example, cell population proliferation (GO:0008283) sub-categories are interspersed amongst the other four top-level GO categories when sorted by correlation strength (Fig. 4B, Supp. Table 1). This lack of a pattern suggests that many more GO processes than might be expected can contribute to cancer. That the GO sub-category, maintenance of cell number (GO:0098727), was amongst the top 10 most correlated GO sub-categories supports our use of cancer as a proxy for cell fitness (Supp. Table 1). If it is found that many, most, or all GO categories (and thus all genes) contribute to cancer, in future studies it will be interesting to classify each into one or more of the categories outlined in the well-known “Hallmarks of cancer” review by Weinberg and Hanahan (31, 32).

PubMed can be considered biased because it is curated by humans and because co-citation does not necessarily tell us the reason for a gene and a disease being mentioned together. Therefore we considered the orthogonal approach of using genome-wide association studies (GWAS) to further characterize the cancer/non-cancer correlations we identified using PubMed. Synthetic interaction testing is a genetic concept, therefore it made sense to consider GWAS because it’s a mainstay for human genetics. We used the UKBioBank dataset because of its broad acceptance and large sample size (46), and specifically the widely-used summary GWAS results of this dataset from the Ben Neale lab (53). We selected a set of 100 cancer and non-cancer conditions (Supp. Table 1) that were most co-cited with genes (as opposed to medical contexts where genes are not reported to be influential). To determine the interaction strength of a gene in cancer vs. non-cancer, we examined the proportion of diseases in each category for which there was an associated SNP in that gene. Our results show a high correlation using either p-value (r = .819, Fig. 4C) or beta value thresholds (r = .812). Because the thresholds we used for p-values (p < 0.001) and beta values (|beta| >= 0.0031) can be considered modest (see Methods for more details) it is possible that the strong correlations could be confounded by gene length. That is, the longer the gene, the more likely it is that it will contain a significant SNP. However, when we control for gene length, our correlations remained statistically significant (p < 0.001). Combined with the results of our PubMed analysis, these findings suggest a strong relationship between the effects of a gene perturbation on cell fitness – represented by cancer – and all other phenotypes – represented by non-cancer conditions.

### Evidence #3: Drug effects on patient cell fitness are reflective of the patient condition response to drug

Like with variation in genes, drugs can have different effects in different genetic backgrounds. We analyzed the published literature to identify citations where the fitness of patient-derived cells in response to a drug was also predictive of the same drug’s effect on the patient’s condition. In this case, *U* is the genetic background of the patient cells and *K* is the drug. We identified 25 studies on PubMed where patient derived cells were generated and treated with drugs and where cell fitness was measured alongside the patients themselves being treated and their drug responses monitored. What was notable in all the studies is that cell fitness was predictive of patient response for widely divergent medications (Fig. 5A). These results might be surprising outside the context of cancer therapy, but they arguably should not be because the level of growth inhibition (inhibitory concentration - IC50) for these drugs varies widely in different cell lines, which have different genetic/epigenetic backgrounds (Fig. 5A, Supp. Table 1).

**Figure 5.**
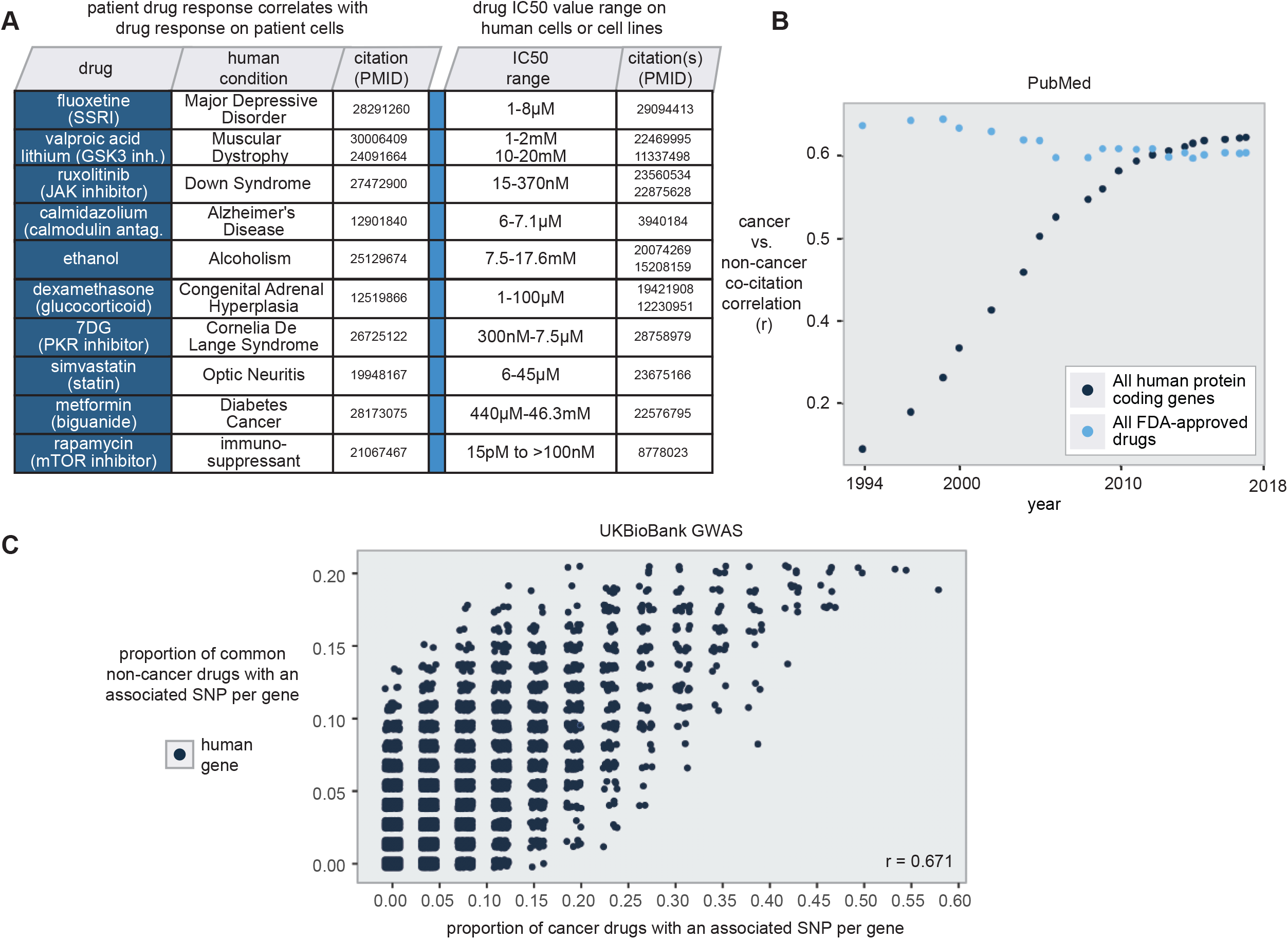
Establishing the efficacy of FDA-approved drugs using cell fitness. **(A)** Drugs where their effects on a patient’s condition and the fitness of the same patient’s cells correlate. Listed drugs met two criteria: 1) Their effect on a patient’s condition and the same patient’s cell fitness were significantly correlated; 2) The drug had differing IC50 values – the concentration at which cell numbers are 50% of their untreated values – in multiple, distinct human cell types or cell lines. **(B)** Correlation (r) over time of human protein-coding genes or FDA-approved drugs co-cited with cancer or non-cancer conditions analyzed as in Fig. 3A. The most recent data point of genes corresponds to the same data that was used in Figure 4A. (**C**) The proportion of non-cancer drugs vs. cancer drugs that have a significant SNP (p-value < .001) in UKBioBank for each protein-coding human gene. These proportions are out of a set of 15 drugs used in UKBioBank patients who had cancer vs. 15 other drugs used in non-cancer conditions that had the highest number of cases in the UKBioBank. Correlation coefficient, r = .671.

Because we found a relatively small number of studies on PubMed that met our strict criteria, we considered the cancer vs. non-cancer paradigm we used in Figure 4 to give us more comprehensive insight on the relationship between drug response and cell fitness. Like with PTEN as previously discussed, our understanding of the role of many genes in cancer has preceded that of our understanding of their roles in other diseases. We reasoned that with several advances, such as various new –omic analyses, our understanding of the molecular basis of many diseases is “catching up” to cancer more recently. Thus, we expect the correlation between all human protein coding genes and all FDA-approved drugs being cited both in and non-cancer to be increasing over time. Indeed, human genes are increasingly being cited with similar frequency in cancer and non-cancer disease contexts (Fig. 5B, Supp. Table 1). This supports the idea that one should be able to increasingly make use of the literature rather than perform experiments to understand an unknown gene’s function. Like with human genes, we found that clinical drugs are commonly cited in both the cancer and non-cancer literature (correlation = 0.599), albeit this correlation hasn’t increased over time like with genes (Fig. 5B, Supp. Table 1). This finding suggests drug repositioning opportunities for non-cancer conditions that are underexplored given the current mindset that identifying drug targets for cancer would require separate efforts from those for non-cancer conditions.

Lastly, to make an analogous analysis with drugs as we did with diseases in Figure 4, we binned the top cancer and non-cancer drugs taken by participants in UKBioBank. Figure 5C shows the proportion of drugs with an associated SNP for both cancer and non-cancer drugs. The cancer vs. non-cancer drug correlation is strong (r = .66) and like with cancer vs. non-cancer diseases in Figure 4 and remains statistically significant when accounting for gene length (Fig. 5C). Combined with the results of our PubMed analysis, these studies lend support to the increased interest in repositioning many non-cancer drugs for cancer (54) (e.g., immunosuppressant, rapamycin and anti-diabetic, metformin), and vice versa (e.g., chemotherapeutics for autism, senolytics for aging (55).

### Proof of concept of the Omniphenotype model using bisphosphonates and its MoA target genes, ATRAID, SLC37A3, and FDPS

We recently published a proof-of-concept of the Omniphenotype model – which can be thought of as an expanded definition of synthetic interaction testing – with the osteoporosis drugs, nitrogen-containing bisphosphonates (43, 56, 57). In this work, we identified a gene network, ATRAID-SLC37A3-FDPS, using cell-fitness-based, drug interaction screening with the widely used osteoporosis drugs, bisphosphonates, and subsequently demonstrated this network controls bisphosphonate responses in cells, in mice, and in humans (43, 56, 57). In this case, bisphosphonates and FDPS were the known, *K,* and ATRAID and SLC37A3 were the unknown, *U,* that we discovered were involved in the bisphosphonates MoA.

In addition to elucidating drug MoA, another strength of the Omniphenotype model is its potential to make new insights into disease biology and therapeutics applications. To take our bisphosphonates findings a step further, we were interested in two questions: 1) How can the bisphosphonates MoA be distinguished from the statins MoA, which also targets the mevalonate pathway but at a different gene, HMGCR, upstream in the pathway (58)? 2) Where in the body might bisphosphonates be therapeutic besides bone? To answer these questions and extrapolate from our cell fitness data to in vivo phenotypes we made use of International Mouse Phenotyping Consortium (IMPC) (59), which to date has performed whole body phenotyping on 6,440 knockout mice.

We were interested to determine how the statins and bisphosphonates MoAs might be distinguished by their in vivo effects. Because ATRAID, SLC37A3, FDPS, and HMGCR deficient mice have not yet been generated by the IMPC, we sought a systematic, well-powered approach that would enable us to indirectly measure the in vivo effects of their loss of function. Using the MCS concept, we wondered to what extent might ATRAID, SLC37A3, FDPS, and HMGCR target genes deficiency, in aggregate, reflect single gene ATRAID, SLC37A3, FDPS, and HMGCR deficiency. By “target genes” we mean genes whose expression levels were affected by ATRAID, SLC37A3, FDPS, and HMGCR deficiency.

To test the validity of this approach we started with bone density because we had an expectation of the sign conventions of the effects we might see based on our previous work and that from others where ATRAID and SLC37A3 inhibition negatively affected bone density and HMGCR and FDPS inhibition positively affected it (57, 60–62). Indeed, based on a comparison of hundreds to thousands of mice deficient in ATRAID, SLC37A3, FDPS, or HMGCR target genes with over 16,000 control mice, ATRAID and SLC37A3 target gene deficiency decreased bone density and FDPS and HMGCR target gene deficiency increased bone density (Fig. 6A).

**Figure 6.**
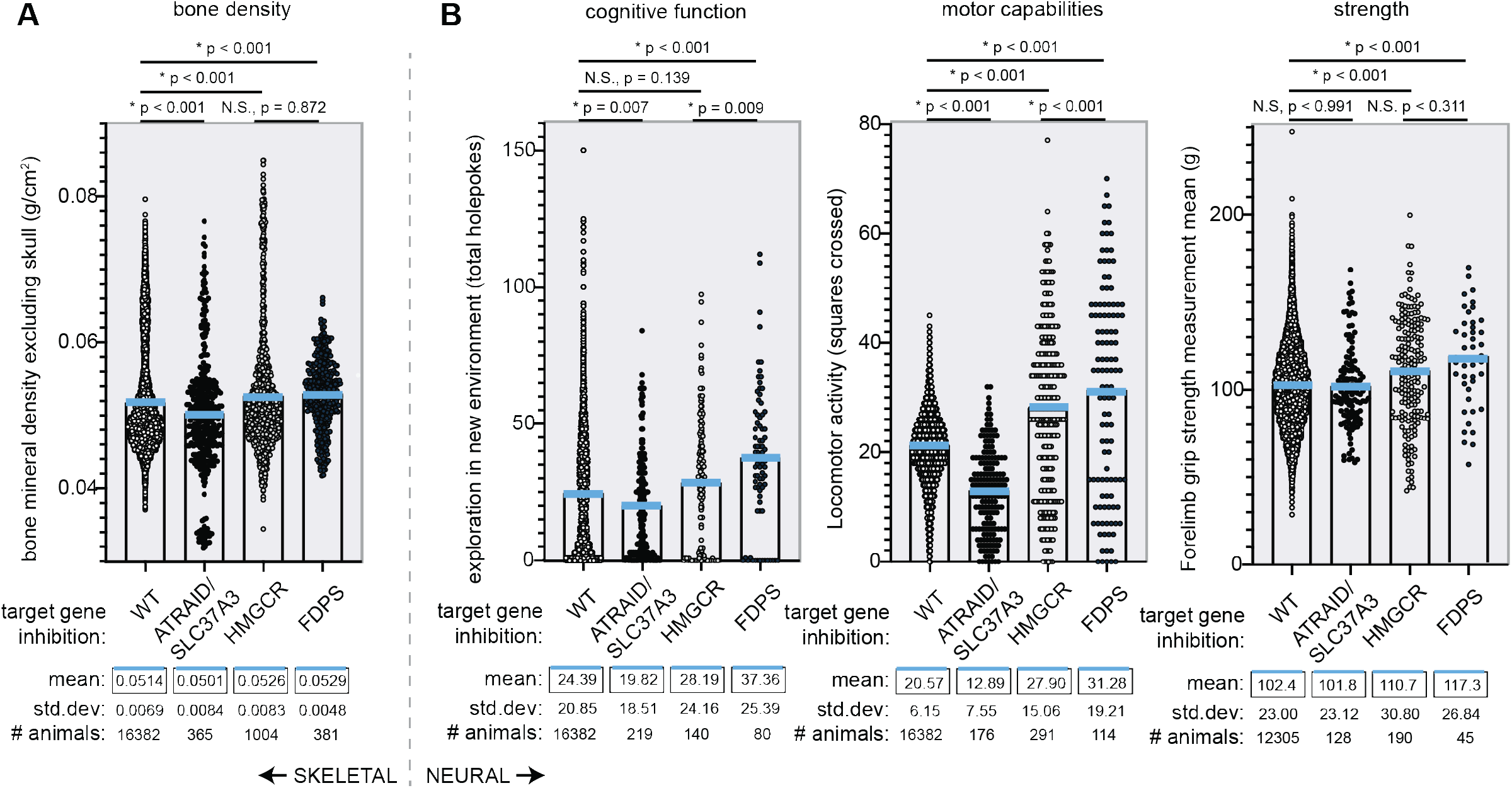
The Omniphenotype model proof of concept using bisphosphonates target genes. In vivo skeletal and neural phenotyping of bisphosphonates MoA target gene output. **(A)** The phenotypes shown are those of aggregated knockout lines of genes of correlating fitness profiles with the indicated bisphosphonates MoA target genes. Descriptive statistics (mean, standard deviation, number of samples) are shown. Red line indicates mean. P-values were obtained using one-way ANOVA corrected for multiple comparisons using the Tukey Method. Number of gene knockout mouse lines considered for each bisphosphonate MoA genotype: bone density | ATRAID/SLC37A3: 23, HMGCR: 66, FDPS, 25. **(B)** Data was analyzed similar to A. Number of gene knockout mouse lines considered for each bisphosphonate MoA genotype: cognitive function | ATRAID/SLC37A3: 23, HMGCR: 9, FDPS, 5. Locomotor Activity | ATRAID/SLC37A3: 11, HMGCR: 18, FDPS, 7. Grip Strength | ATRAID/SLC37A3: 8, HMGCR: 12, FDPS, 3.

Like with bone density, our cell fitness data predicted that ATRAID and SLC37A3 inhibition should negatively regulate neurological behaviors and FDPS inhibition should positively regulate it. Moreover, FDPS inhibition should have stronger effects than HMGCR inhibition. Consistent with these predictions, ATRAID and SLC37A3 target gene deficient mice had decreased cognitive function and motor activities (Fig. 6B). For both activity tests, the beneficial effects of FDPS target gene deficiency were statistically significantly greater than those of HMGCR target gene deficiency (Fig. 6A, B). For the other phenotypes including bone density, FDPS target gene deficiency trended to being more beneficial than HMGCR target gene deficiency. These results provide a preliminary indication that ATRAID and SLC37A3 inhibition might promote neurodegenerative symptoms and FDPS inhibition, e.g., with bisphosphonates treatment, would potentially alleviate them and do so more than HMGCR inhibition.

## DISCUSSION

We propose cell fitness as a simple, yet comprehensive approach to rank less-studied genes, cell processes, diseases, and therapies in importance relative to better-studied ones (Fig. 7). Rather than needing ever more refined experiments, this work suggests the antithetical approach. That is, one can use fitness in mammalian cells to estimate the importance of a poorly characterized factor of interest (herein referred to as *U*) in a biological context of interest by comparing the new factor’s individual effect on cell fitness vs. its combined with a factor with a known role in the biological context of interest (herein referred to as *K*) (Fig. 7). Besides the bisphosphonates and the statins, there are numerous other examples in the literature such as with the mTOR pathway and rapamycin (63) where the steps from cell fitness to phenotypes relevant in people can be readily traced.

**Figure 7.**
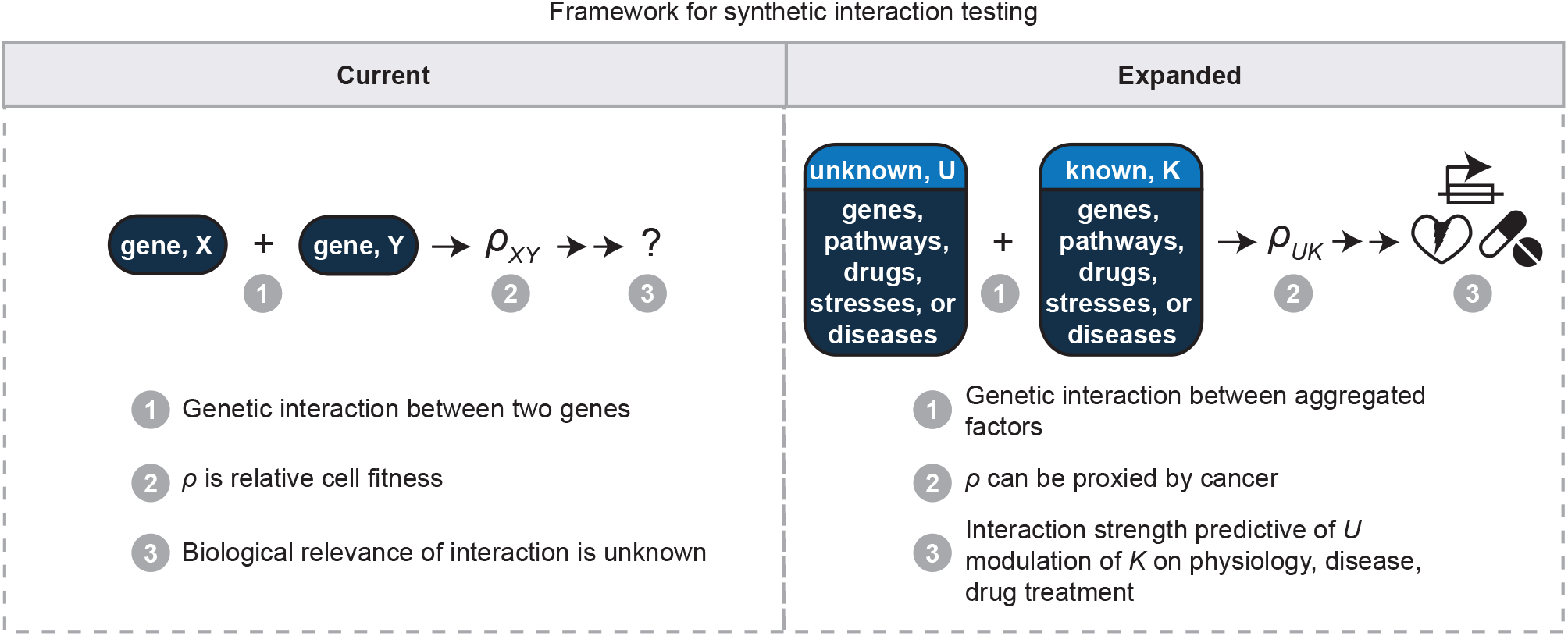
Current vs. expanded conceptual frameworks for synthetic interaction testing. A currently, commonly used framework for synthetic interaction testing is as follows: 1) Genetic interactions are tested between two genes, e.g., *X* and *Y*; 2) *ρ,* the phenotype being measured, is relative cell fitness; 3) It isn’t known how the interaction strength between *X* and *Y* might be relevant to physiology or disease. The expanded framework described herein is as follows: 1) Genetic interactions can be tested between aggregated factors such as multiple genes and/or pathways (related to Figure 3); 2) *ρ* can be proxied by the relative involvement of the interactors in cancer (related to Figure 4); 3) One interactor, *K*, has a known effect on a physiology or disease of interest. The effect of the other, *U*, in that context can therefore be measured relative to *K* (related to Figure 5). The implications of mammalian cell fitness-based synthetic interaction testing as described here is that the results using cell fitness can be readily extrapolated to the ultimate phenotype of interest, e.g., osteoclast resorptive ability. That is, the magnitude of the synergy on cell fitness of the two factors under consideration is expected to be proportional to the magnitude of their synergy on the phenotype of interest.

Cell fitness is a phenotype that has long had traction in the yeast and bacteria communities, but not yet in mammalian systems outside of cancer. To increase awareness of its utility, below we address its relevance as a phenotype to the different levels at which mammalian biologists tackle their work: at the level of cell and molecular processes, at the level of disease, and at the level of environmental factors such as drugs.

### Relevance to cell and molecular processes (Fig. 3)

Our Figure 3 results suggest one can study the majority of cell and molecular processes using cell fitness-based, synthetic interaction testing. Specifically, Figure 3 shows that to the extent that cell and molecular processes are well represented by GO terms, essential genes as determined by cell fitness (cell counting) are involved in the vast majority of such processes. This is useful information for molecular biologists because we often spend considerable time developing experimental models. For example, large efforts are often spent upfront (before getting to the question of interest) developing technology, such as cell types to model various tissues. This is done because of the often-unquestioned assumption that the cell type being studied matters. While this can be true, it is worth considering whether it’s the rule or the exception. In support of the latter, in single cell RNAseq analysis, 60% of all genes are expressed in single cells and the percentage jumps to 90% when considering 50 cells of the same type (64). Similar percentages are obtained at the population level when comparing cell types (65, 66). This argues that the majority of the time, there should be a generic cell context to do at least initial “litmus” synthetic interaction testing of a hypothesis concerning a new factor of interest, *U,* before proceeding with a more complex set-up. Like with cellular phenotypes, many molecular phenotypes, such as reporter assays require domain expertise, can be hard to elicit, and might not be easily reproducible or scalable. Whereas cell fitness is robust and readily approximated by inexpensive assays, such as those to measure ATP levels.

### Relevance to diseases (Fig. 4)

Our PubMed and GWAS analyses lend growing support to using dry-lab approaches as first-class tools for discovering medically relevant biology. The costs to perform the wet-lab experiments required to publish in the “-omic” era are not small and favor well-resourced teams. Gleaning insights from publicly available data, which is large and growing rapidly, and accessible by anyone with programming skills, is a way to level the playing field.

One counterintuitive insight from Figure 4 is that core genes on the diagonal might not be as good drug targets as genes off the diagonal because the core genes are involved in more diseases. This might mean that because of their importance, core genes might be too networked with other genes to be targeted by therapies without the therapies causing off-target effects. Fortunately, this might then also mean those genes off the diagonal might make more appealing drug targets because they are less networked. Thus, in this case we would use cell fitness as a filter to prioritize genes for further study. This is important because like with income inequality where the “rich get richer” certain genes get cited over and over (44), and approaches to aggregate data are known to favor the “discovery” of already well-studied genes over less studied genes (67). When applied in unbiased methods such as with genome-wide CRISPR-based screening (43), we expect our approach to be particularly impactful in addressing this issue of identifying important genes that historically are ignored.

### Relevance to environmental factors, e.g., drugs (Fig. 5)

With the exception of oncology, it could be argued pharmacogenomics is only slowly going mainstream in many fields. One reason for this is the poor feasibility and high cost-to-benefit ratio of current genetic testing strategies to predict drug response (68). Though more work needs to be done to test the generalizability and “real-world” application of the studies we highlight in Figure 5A, they do provide important initial evidence for exploring cell fitness as a diagnostic assay for precision medicine. In addition to drug diagnostics, these studies have implications for drug discovery. Both target-based and phenotypic-based drug discovery have drawbacks (69). Target-based approaches don’t necessarily tell you about the phenotype, and phenotypic screens don’t tell you about the target. On the other hand, cell fitness-based, drug response genetic screening (70) can potentially be the best of both worlds because it extrapolates well to the phenotype of interest and provides the gene target in the same screen.

Still, we acknowledge limitations to our findings. Our work focuses on genes and drugs as the genetic and environmental variables of interest. Future work must determine to what extent our work would apply to other variables, such as epigenetic signatures. PubMed-based analyses are subject to biases. Many genes are not published on due to human factors. Even among those genes that are published, their representation is affected by publication bias towards positive results. Additionally, our citation analysis relies on curation by NCBI, which due to its high stringency and the current limitations of natural language processing strategies, likely leaves out a large number of studies that should be included. However, this is why we also analyzed GWAS, which provides a more unbiased look. As more methods get developed and more knowledge accumulates, a scientist’s job of figuring out where to focus their attention gets harder. Our view is that cell fitness could be used more often to help scientists triage their opportunities.

## FIGURE LEGENDS

**Figure S1.**
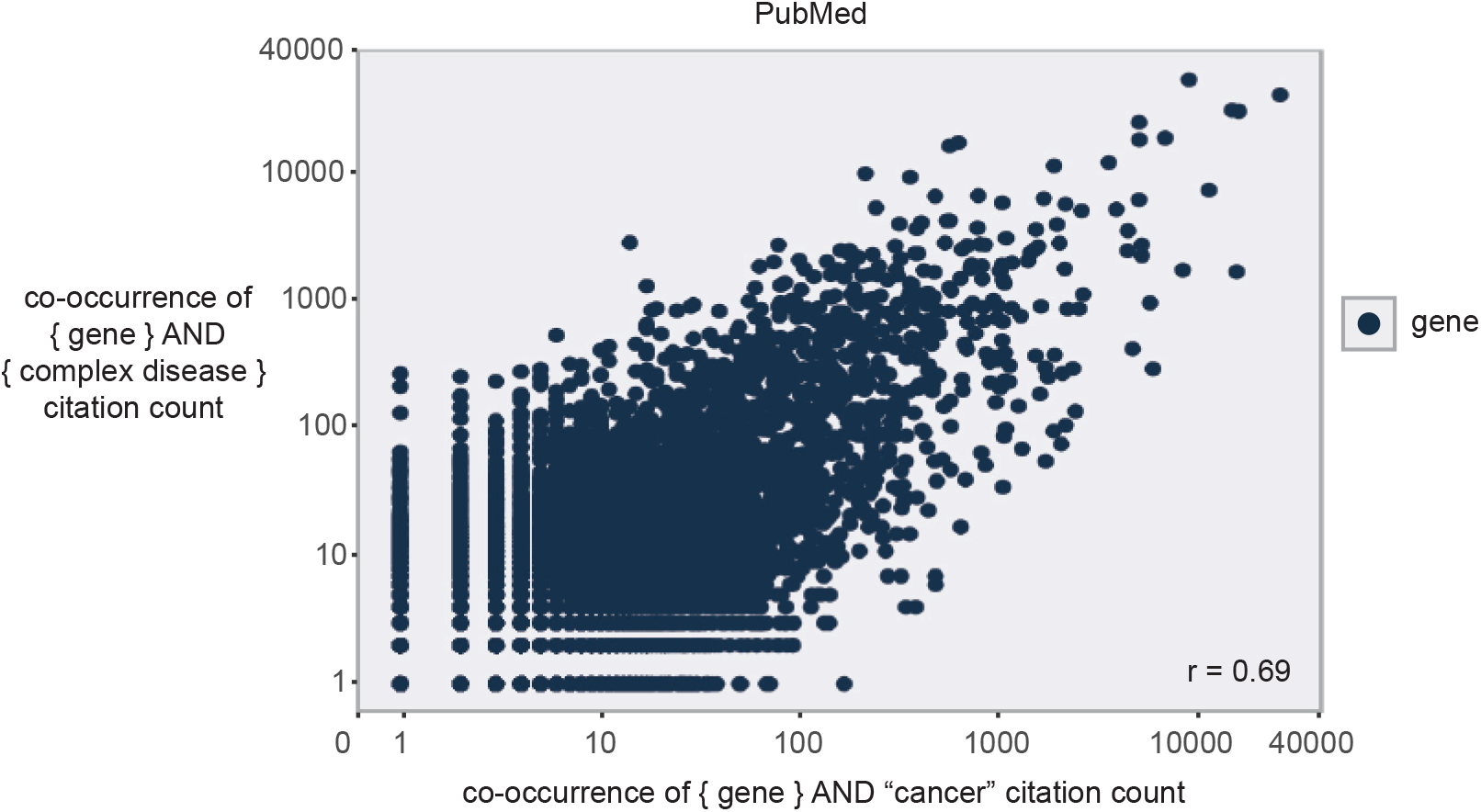
Common disease-restricted cancer vs. non-cancer PubMed analysis. Nearly all common disease genes are cancer genes. The co-occurrences of each human gene with cancer (x-axis) and non-cancer (y-axis) conditions in ~27 million PubMed citations were counted. The following non-cancer conditions were considered: infection, Alzheimer’s, cardiovascular disease, diabetes, obesity, depression, inflammation, osteoporosis, hypertension, stroke, and inflammation. Line of best fit - Slope, 0.75; Correlation coefficient, r = 0.69.

**Table S1 | Data related to all figures.** Data to generate scatter plots as well as list of diseases and drugs analyzed are included in Supp. Table 1.

## AUTHOR CONTRIBUTIONS

T.R.P, N.C.J, and J.P conceptualized the project. N.C.J and T.R.P performed the analysis. N.L.S. provided feedback on the GWAS analysis. T.R.P wrote the paper and N.L.S, N.C.J, and J.P edited it.

## METHODS

### Code

All code created for this work is available at: https://github.com/tim-peterson/omniphenotype. All unreported p-values for correlation coefficients are to be assumed to be so small that R rounds them down to 0. Our analyses throughout rely on gene symbol identification. We used NCBI official gene symbols in all cases to obtain data for each gene. The biases using this approach penalize against gene symbols that use common English words, such as “MICE”, or that are short in length, e.g., less than 4 characters. These criteria do not bias for any particular biological function, therefore they appeared to us to be a reasonable compromise between being comprehensive with our analysis while still maintaining a feasible workload.

### Budget analysis (Fig. 1A)

A list of high-confidence gene interaction pairs was created by incorporating protein-protein interaction and co-dependency, a.k.a., co-essentiality or imputed synthetic interaction, analyses. Protein-protein interaction data was obtained from BioPlex at https://bioplex.hms.harvard.edu/interactions.php (34, 71). Interactions detected in both HEK293T and HCT116 cell lines comprised our high-confidence protein-protein interactions list of 21,729 interactions. Imputed synthetic interaction analyses were performed by identifying the top 100 correlating genes in the DepMap with the list of 21,729 (40).This resulted in a list of 5,824 interactions, which represent ~27% of the total. To calculate the cost of studying these pairs, we used the NIH grant funding database, Federal RePorter (73). Abstracts and project terms for NIH-funded grants were downloaded by year from 2000 to 2020. These datasets were queried to obtain grants that contained the names of both genes in each pair (of the 5,824) and the funding amount (in US dollars) for that grant. The cumulative number of studied pairs was taken for each year and extrapolated using linear regression to find the date and cumulative dollar amount when all pairs would be studied. The dollar amounts were multiplied by how much is estimated to be needed to convert this funding to FDA approved drugs, i.e., 44,200/670 = 66X (35). From 2000 to 2020, $44,200 million is estimated to have been needed in total private investment to result in the approval of 18 drugs where $670 million in total public funding was used to develop the science that led to those 18 drugs.

### Gene ontology and cell essential gene analysis (Fig. 3A)

A list of all human genes was obtained from NCBI, ftp://ftp.ncbi.nih.gov/gene/DATA/GENE_INFO/Mammalia/. Gene ontology (GO) information for each human gene was obtained from BioMart, https://useast.ensembl.org/info/data/index.html. By “essential”, we refer to cell essential – required for viability of a cell – as opposed to organism essential – required for viability of an organism. Cell essential gene lists were obtained through CRISPR screens performed by Hart et al. and Wang et al. and haploid screens performed by Blomen et al (45–47). If knockdown of a gene resulted in a statistically significantly decreased cell count (p < .05 or FDR of 5%) then the gene was determined to be cell essential.(45–47). There were ten cell lines tested: K562, KBM7, Jiyoye CS, Raji CS, HAP1, HCT116, DLD1, HeLa, GBM, RPE1. For the relaxed filter data point, we used cell-count based fitness scores at p < 0.1 for both the Wang and Hart studies. For the Blomen study, p-values > 0.05 weren’t given, therefore we took all genes with fitness scores less than 0.5 standard deviations above the mean.

### Gene inactivation fitness profile correlation (Fig. 3B)

Single gene inactivation cell count-based cell fitness profiles from the Broad Institute and Sanger Institute were downloaded from the DepMap website, https://depmap.org/portal/download/ using the 2019q2 dataset. Fitness profiles for single genes from the same GO category, which were chosen because they are highly co-cited. Pair-wise Pearson correlations were then calculated using the python pearsonr() function. P-values were false discovery rate (FDR) corrected using Benjamini and Hochberg methods. In python, this is set using SciPy as shown here: https://github.com/tim-peterson/omniphenotype/blob/master/figure3B.py#L96. Co-citation gene-gene cell fitness enrichment analysis was performed by counting all genes in the top 20 with greater than one citation compared with the total number of genes with greater than one citation.

### PubMed analysis (Fig. 4 and 5)

In both Fig. 4 and 5, PubMed was programmatically accessed using the E-utilities (https://www.ncbi.nlm.nih.gov/books/NBK25497/) API endpoint: https://eutils.ncbi.nlm.nih.gov/entrez/eutils/esearch.fcgi. From this endpoint, PubMed IDs (PMIDs), were counted (Fig. 4A-B, 5B), or abstracts were analyzed for the relevant information (Fig. 5A).

#### Cancer vs. non-cancer analysis (Fig. 4A)

All PMIDs for each human gene and their homologs and all PMIDs for each Medical Subject Headings (MeSH, https://www.nlm.nih.gov/mesh/meshhome.html) were collected into tab-delineated files. A citation was determined to be associated with a particular MeSH term if it came up from searching PubMed with “<MESH term> [mesh]”. A citation was determined to be associated with a particular gene if it has been labeled in “Related articles in PubMed” within the Gene resource of NCBI. See Omniphenotype Github repository for how these application programming interface (API) calls were made. The gene-PMID file was intersected with the MeSH-PMID file, and the resulting gene-MeSH term list was separated by the MeSH term “neoplasm”, which incorporates the terms “cancer”, “malignancy”, “tumors”, and other neoplasm-related terminology. Meaning, all citations that were annotated as referring to a gene that were co-annotated with the MeSH term “neoplasm” were considered a “cancer” citation and all other citations that were co-annotated as referring to a gene and any other disease MeSH term besides “neoplasm” were considered “non-cancer” citations. Those citations which mapped to both neoplasm as well as other conditions were considered “cancer” citations. Genes with less than 10 citations were excluded from the analysis. The correlation co-efficient of cancer vs. non-cancer citation counts for all genes was calculated using R using the lm() function on log-log data.

#### Gene Ontology-subsetted, cancer vs. non-cancer citation analysis (Fig. 4B)

To determine the diversity of correlations for cancer vs. other conditions among GO categories, we took a subset of major GO categories (immune system process, metabolic process, behavior, developmental process, and cell population proliferation) as a representation of a diverse subsampling of biological processes. We then iterated through the sub-category/child terms of each of these major GO categories and found the correlation of all cancer vs. non-cancer citations for genes associated with that term. The results are displayed as a 1-Dimensional heatmap sorted by correlation strength with a secondary mapping of the higher order GO categories also displayed.

#### GWAS Analysis (Fig. 4C and 5C)

Beta-values and p-values for each SNP-disease and SNP-drug pairing were generated using the Ben Neale lab analysis of UKBiobank data: http://www.nealelab.is/uk-biobank. SNPs were assigned to protein-coding genes according to the GENCODE release 19 (GRCh37.p13) by specifying if a SNP overlapped any portion of a gene (first to last exon), then it pertained to that gene. This means that a SNP could be assigned to multiple genes. The proportion of conditions with an associated SNP was obtained by finding the proportion of cancer and non-cancer conditions that had any associated SNP for each gene. The phrase “associated SNP” as used throughout refers to the following thresholds: a p-value less than 0.001 (10^-3^) and a beta-value greater than 0.0031. The logic for these thresholds derive from the omnigenic model which posits that a large number of genetic variants with small effect size influence each phenotype (31). This model has been borne out in many studies including those computing polygenic risk scores, which assess the contributions of many DNA variants to the heritability of complex diseases and traits (74–76). For example, height is influenced by a large number of variants with small effect size (77, 78). Consistent with the omnigenic model, using traditional GWAS thresholds such as 10^-8^ for p-values we found few genes that met this threshold per UKBiobank dataset, which precluded our analysis of cancer vs. non-cancer correlations. To ensure robust results, we tested a range of p-values and beta-values and established 10^-3^ and 0.0031, respectively, as producing high gene coverage while still being stringent compared to thresholds such as p-value < 0.05 that are generally accepted with smaller datasets.

#### Disease Selection (Fig. 4C)

A set of the top 100 diseases were selected by ranking MeSH terms in the Diseases category by how often they were co-cited with genes in PubMed. This produced a list of diseases that were the most commonly studied in molecular biology contexts, i.e., in the context of genes. Within this ranking, we used the datasets corresponding to the self-reported illness codes in UKBiobank. We then separated these diseases into cancer and non-cancer diseases.

#### Drug response in patients vs. patient cells analysis (Fig. 5A)

To identify studies that reported findings on patient cell drug responses that correlated with the patient we used two search terms that are commonly associated with culturing patient cells: “human lymphoblastoid cell lines” and “peripheral mononuclear blood cell”. The phrase “human lymphoblastoid cell lines proliferation” was queried in PubMed and 804 abstracts were returned. “PMBC” as in peripheral mononuclear blood cells was substituted for lymphoblastoid cell lines (LCLs) to identify the simvastatin study. The LCL and PMBC citations were manually curated to identify relevant citations that mentioned drug responses on cell proliferation. The IC50 studies were obtained by querying [drug_name] AND “cell viability” OR “IC50”.

#### Co-citation time series analysis (Fig. 5B)

PubMed PMIDs are largely chronological, i.e., higher PMIDs mostly mean a more recent citation and vice versa, with the exception of PMIDs < 8M as well as those between 12M and 15M. As of July 9, 2019, citations up to the end of 2017 have been annotated with MeSH terms and gene associations such that cancer vs. non-cancer correlations for multiple time points could be determined. Citations were split every million PMIDs and for the purposes of graphing the results the calendar year was approximated. The FDA-approved drug list was obtained from: https://www.fda.gov/drugs/drug-approvals-and-databases/drugsfda-data-files. Similar to the cancer vs. non-cancer analysis for genes, all PMIDs for each drug were collected into a tab-delineated file, intersected with the MeSH-PMID file, and the resulting list was separated by those citations that were co-annotated with the MeSH term “neoplasm” and those that were co-annotated with other disease MeSH terms.

#### Drug Selection (Fig. 5C)

Drugs were defined as cancer-associated if greater than 10% of their PubMed citations were co-cited with the MeSH term “Neoplasms”. Of the drug codes in UKBiobank categorized in the Neale lab dataset, 15 fit this categorization. An additional 15 drugs were chosen in non-cancer-associated category by ranking drugs by the number of cases in UKBiobank. The names of each drug are listed in Supp. Table 1.

### Bisphosphonate proof of concept (Fig. 6)

The inclusion criteria for a gene knockout mouse line to be included in the analysis was as follows: ATRAID or SLC37A3 deficient cells showed < 0.75 expression levels of the gene relative to wild-type and < 1 for both ATRAID and SLC37A3. HMGCR or FDPS inhibited cells showed > 1.5-fold increased expression of the gene relative to wild-type, which means the phenotyping data for these knockout lines had to be ‘inverted’ by normalizing to wild-type. Gene expression from a microglia proxy (K562) cells was used, but data from a neuronal proxy (HEK293T) showed a similar pattern (data not shown). Phenotyping data is from the International Mouse Phenotyping Consortium (IMPC).

## Supporting information

Omniphenotype_supp_table1

